# The control of targeted jumps in nymphal praying mantises

**DOI:** 10.64898/2026.04.30.721899

**Authors:** G. S. Girish Kumar, Sanjay P. Sane

## Abstract

Arboreal insects have developed various strategies to navigate their discontinuous habitats. Many insects, including leafhoppers, katydids, and praying mantises, exhibit the ability to actively leap across their leafy platforms and land on a distant substrate. This behavior is especially important for non-winged insects, including nymphal forms of winged insects, which cannot fly between these substrates. To make a targeted jump, an animal must first orient towards the target, estimate the target distance and angular location, and jump with the appropriate take-off speeds and angles to land on their intended substrate. In three-dimensional space, jumping from one point to another requires estimating distance, as well as azimuthal and elevational angles. Jumping insects such as mantises typically reorient their bodies on the substrate to align with the azimuthal direction of the target. This behavior effectively reduces the task to a two-dimensional problem, in which they must estimate only the distance to the target and its elevational angle. Many insects, including praying mantises, perform rhythmic lateral head movements called ‘peering’ before performing a targeted jump. Although previous studies suggest that mechanisms such as motion parallax while peering are used for distance estimation, the full repertoire of behaviors that enable mantises to jump to arbitrarily located substrates remains unclear. We hypothesized that mantises have distinct behaviors for distance and elevation angle estimation, which enable them to independently modulate their take-off speeds and angles before jumping. To test this hypothesis, we developed behavioral assays in which mantises were placed on a launch platform and jumped to a target platform positioned at variable distances and angles. Using this apparatus, we filmed the jumps of Giant Asian mantis nymphs (*Hierodula* spp.) with high-speed videography and tracked body parts to quantify take-off speed and angle. Because mantis jumps are ballistic, their trajectories can be modeled as projectile motion. Our results indicate that mantises estimate target distance and elevation angle using two separate behavioral strategies: distance is assessed through peering maneuvers that generate motion parallax, whereas elevation angle is determined through visual fixation of the target accompanied by specific postural adjustments. By combining these behaviors, mantises modulate the magnitude and direction of propulsive force to achieve successful jumps.

## Introduction

For arboreal insects, the ability to jump across gaps from substrates of varying texture and compliance is key to their locomotor success. In wingless insects, including the nymphal stages of many winged insects, jumping is largely a unitary event; once airborne, their center of mass follows the parabolic curve of a typical projectile. This means that before jumping, insects must estimate the distance between the substrates and generate *a priori* the force with which to push on their substrates. In these cases, the insect adopts a stationary posture before taking off; such jumps are therefore termed standing jumps (Alexander, 2013). The question of how insects gauge the variable gaps between substrates and control their jumps is thus central to our understanding of arboreal locomotion.

Insects jump in a wide range of behavioral contexts, including initiation of flight, escape from threats, catching prey, or navigating across discontinuous surfaces. Flight initiation in many adult insects involves a jump accompanied by wing flapping, either in sequence or simultaneously, to become airborne (Brackenbury, 1996; Trimarchi and Schneiderman, 1995a; Burrows and Dorosenko, 2015). Jumping is also essential for many juvenile hemimetabolous insects, which do not possess wings, and most wing-bearing adult insects to escape threats (Tauber and Camhi, 1995; Trimarchi and Schneiderman, 1995a; Santer et al., 2005). In these forms of jumping, the landing substrate may not be pre-determined by the insect, and the insect primarily intends to escape from its current location. However, in many contexts, wingless forms of insects, such as locusts (Sobel, 1990a), mantises (Walcher and Kral, 1994), and adult winged insects including flea beetles (Brackenbury and Wang, 1995), leafhoppers, froghoppers, and spittle bugs (Brackenbury, 1996) perform targeted jumps. These insects occupy a wide range of habitats and can jump from substrates with markedly different mechanical properties. They launch from hard surfaces such as rocks, from smooth and compliant substrates such as leaves seen in arboreal taxa including leafhoppers (Clemente et al., 2017), froghoppers (Goetzke et al., 2019), locusts (Taylor et al., 2024), and mantises (Goetzke et al., 2025), and even from water surfaces, as in pygmy mole crickets (Burrows & Picker, 2010), water striders (Hu and Bush, 2010), and long-legged flies (Burrows, 2013). Correspondingly, many species exhibit habitat-specific morphological adaptations of the legs that enhance traction while jumping: adhesive pads facilitate attachment to smooth substrates in leafhoppers (Clemente et al., 2017) and mantises (Goetzke et al., 2025), spines and hooks improve grip on rough or soft surfaces in froghoppers (Goetzke et al., 2019), and specialized structures such as tibial paddles and spurs in pygmy mole cricket (Burrows and Picker, 2010) and hydrophobic hairs enable effective interaction with water in water striders (Hu and Bush, 2010). In natural settings, and especially in arboreal environments, both gap distances and substrate mechanics vary over orders of magnitude (Gilman and Irschick, 2013), placing substantial demands on locomotor performance. The phylogenetic and ecological diversity of these taxa has thus led to a wide range of morphological and behavioral adaptations for jumping, making it a compelling system for studying locomotion from both biomechanical and neurobiological perspectives.

Before executing a jump, insects orient toward the target, estimate its distance and elevation, and generate appropriate take-off speed and angle to reach it. Goal-oriented behaviors such as targeted jumps thus require sensorimotor integration and precise motor control. These processes are best studied in systems in which insects reliably jump toward specific, well-defined targets. Praying mantises are mostly diurnal, arboreal predatory insects which rely heavily on visual feedback during prey capture and possess a fovea with high visual acuity (Barros-Pita and Maldonado, 1970; Maldonado and Barros-Pita, 1970; Rossel, 1980). They have binocular vision, which enables distance estimation through stereopsis (Rossel, 1983; Nityananda et al., 2016). In addition to stereopsis, they can also estimate distance using motion parallax, through a behavior known as peering, in which the body oscillates sideways relative to the target (Horridge, 1986; Walcher and Kral, 1994; Poteser and Kral, 1995). During a jump, they perform controlled aerial maneuvers and landings by exchanging angular momentum within their body parts during a jump and the take-off speeds scale with body size (Burrows et al., 2015; Sutton et al., 2016). In addition to distance, mantises must also estimate the angular position of the target along the azimuthal and elevational axes. It remains unclear, however, whether these two spherical coordinates are computed through shared sensorimotor pathways or via distinct, partially segregated circuits. Of these two angles, mantises can adjust the azimuthal component by turning towards the target, thereby reducing the problem to just estimating the elevational angle. Once distance and elevational angle are determined, mantises can generate the appropriate take-off force in the relevant elevation to reach the target.

We hypothesized that mantises modulate both their take-off speed and take-off angle based on a separate estimation of target distance and target angle. To test this hypothesis, we developed two behavioral assays in which mantises were placed on a launch platform and motivated to jump to a target platform that was placed at adjustable distances or at adjustable angles (in elevation). Our data show that mantises assess target distance using peering and the elevational angle of the target through visual fixation, followed by body positioning. Feedback generated from these two behaviors thus determines the motor planning for the jump.

## Methods

### Animal procurement

We collected nymphs of the giant Asian mantis (*Hierodula* spp.) at various developmental stages from the gardens of the National Center for Biological Sciences (NCBS), Bangalore, India. Because our sampling was opportunistic, we collected individuals spanning developmental stages from first instar to final instar. Across these stages, body mass ranged from 0.02 - 1.925 g (~100x difference), and body length ranged from 16.31 - 70.85 mm (~ 4x difference; see Supplementary Tables 1, 3, 5). We used the captured nymphs for experiments within two days of collection and weighed them before testing. To isolate the contribution of the legs to jumping, we only worked with wingless nymphs in all experiments. The nymphs could only be identified to the genus level, because morphological identification keys exclusively focus on the adult form in the genus *Hierodula*. After completing the experiments, we preserved each nymph in 70% ethanol for future reference.

### Experimental conditions and filming

We filmed all jumps at 2000 Hz with an exposure time of 0.2 ms and a resolution of 1280 × 720 using a high-speed camera (Phantom VEO 640L, Vision Research Inc, Wayne, New Jersey, USA) fitted with a 24–85 mm lens (AF-S NIKKOR 24–85 mm f/3.5–4.5G ED VR, Nikon Corporation, Tokyo, Japan). To illuminate the filming region uniformly, we used a 30 W LED with a diffuse reflector, which provided a maximum light intensity of 7000 lux (measured with a Center 337 Light Meter, Center Technology Corp., New Taipei City, Taiwan). Because the jumps occurred largely within a plane, a single orthogonally placed camera was sufficient to capture the key features of each jump. We attached a 30 mm printed scale to the setup to calibrate the videos and convert pixel measurements to metric units. We captured videos using the Phantom Camera Control (Vision Research Inc, Wayne, New Jersey, USA) application and saved them in the .cine format. We then exported the videos at a playback speed of 25 fps as .avi files with frame number, image resolution, frame rate, and exposure time printed at the bottom of each frame. Finally, we converted the videos to .mp4 format using FFmpeg (FFmpeg developers) to reduce file size and enable further analysis. The ambient room temperatures in all these trials ranged from 25 to 28 °C.

### Setup to investigate the effect of target distance on take-off speed and angle

The setup consisted of two posts of equal height attached to a linear slide rail, allowing the distance between them to be adjusted (Fig. 1A). Each post was capped with a horizontal foam platform. In all the experiments, we placed the mantis on one platform (hereafter termed the *take-off platform*) and induced it to jump to the other platform (*target/landing platform)* by manually moving a black rod or pencil positioned farther than the target platform to attract the attention of the mantis. Following this step, the mantis consistently performed lateral body movements (*peering behavior*) before jumping. Mantises typically peered first at the rod and then at the edge of the target platform, after which they jumped. If a jump took longer to elicit, we gently directed a stream of air from behind the mantis to induce it to peer and jump. The minimum distance between the platforms was empirically determined by adjusting the distance until the mantis needed to jump to reach the other side, rather than reaching over with its long legs. The target distance (edge to edge) was increased in increments of either 10 mm or 20 mm steps until the mantis, rather than jumping, turned away from the target after peering. Each individual underwent approximately five trials per target distance, with the number of distances varying across individuals depending on body size. Using this setup, mantises could be induced to jump repeatedly, underscoring their readiness to jump. We did not observe any signs of adaptation or fatigue over the course of an experiment.

**Figure 1.**
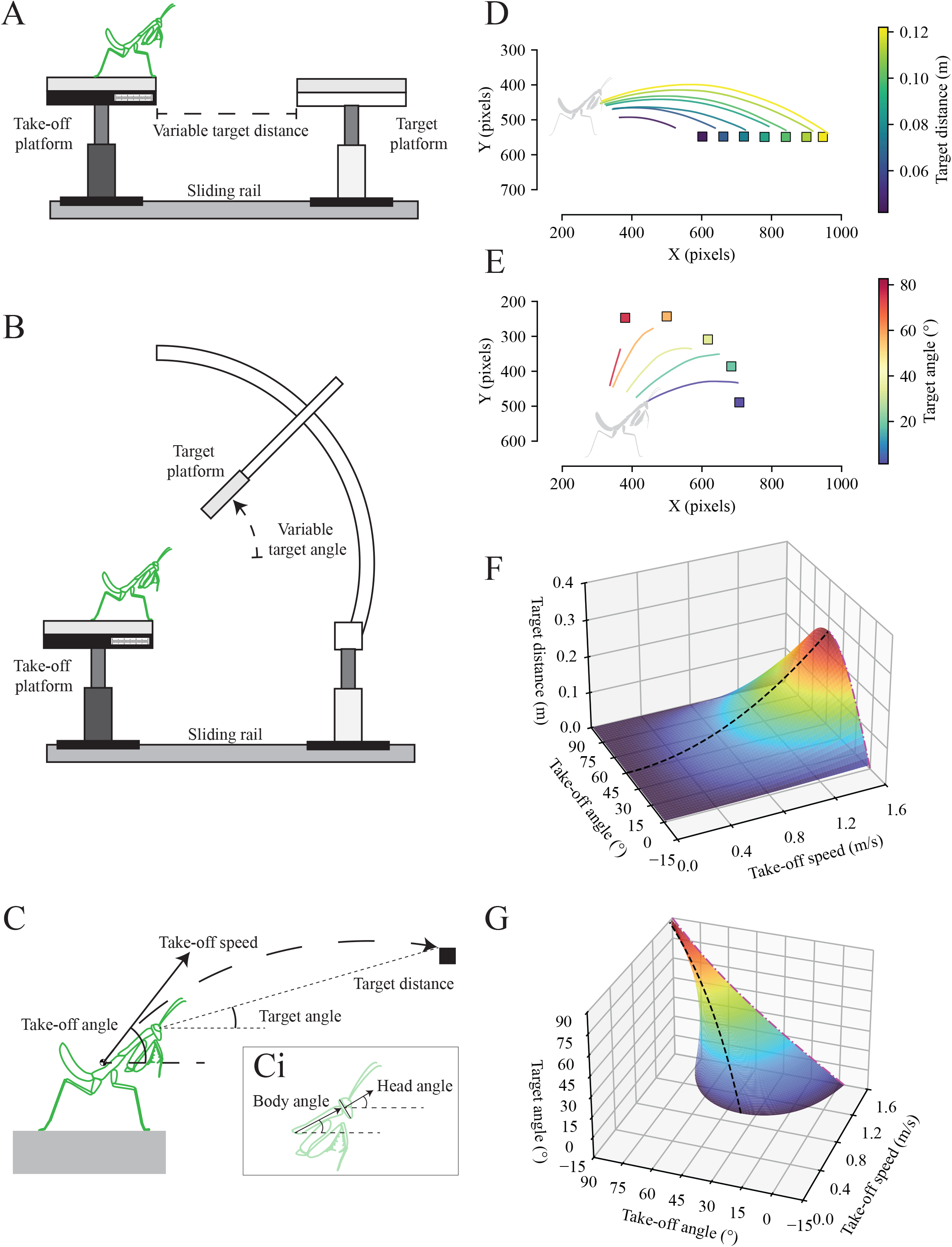
Methodology used to investigate the control of jump in mantises. (A) Experimental setup for varying target distance. The setup consists of two posts mounted on a sliding rail with foam platforms, allowing the distance between them to be adjusted. (B) Experimental setup for varying target angle. The setup is similar to (A), with the target platform attached to a rod slotted into a 3D printed arc centerd on the eye of the mantis, enabling controlled variation of target angle. (C) Schematic illustration of both setups showing the key measured parameters: take-off speed, take-off angle, target distance and target angle. (Ci) Inset showing body angle and head angle. Representative CoM trajectories from an individual mantis in (D) the varying target distances experiment and (E) the varying target angle experiment. (F-G) 3D state-space of projectile motion where (F) the target angle is fixed to 0° and (G) the target distance is fixed at 0.12 m. With the take-off speed capped to 1.6 ms^−1^ in the model, the black dashed line indicates the *speed/energy minimizing strategy* and the magenta dot & dashed line refers to the *time minimizing strategy*.

### Setup to investigate the effect of target angle on take-off speed and take-off angle

To alter the target angle of jumps in elevation, we included a take-off platform similar to the one described above, but equipped with a 3D-printed arc assembly that held a rod with a foam platform attached at the distal end (Fig. 1B). This platform could then be positioned from 0° to 90° in 22.5° increments. We centerd the arc on the mantis’s eyes, allowing the angle of the foam platform to be varied in elevation while keeping its distance (~1.5 body lengths) to the mantis constant. Each individual underwent approximately five trials per target angle.

### Digitization and quantification of take-off speed and angle

Ignoring drag during the aerial phase of the jump, the jump trajectory is expected to be parabolic, following the properties of projectile motion. Thus, the jump trajectory can be defined by its take-off speed and take-off angle by approximating the mantis’ body to a point particle positioned at the center of mass (CoM). Because the body contorted during the jump, the position of the CoM could sometimes lie outside the body, making it harder to track the CoM using any specific body features. To circumvent this, we subdivided the mantis body into rigid body parts whose CoMs could be assumed to be the same as their geometric centers (Supplementary Fig. 1). The CoM of the body (X_CoM,_ Y_CoM_) was obtained using the following equations:

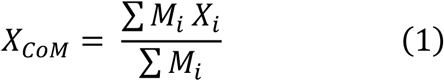

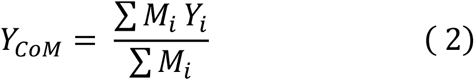

where *M*_*i*_ is the mass of the i^th^ segment, and *X*_*i*_ and *Y*_*i*_ are the X- and Y-positions, respectively. Following the method described in Burrows et al. (2016), we measured the mass of each body part after euthanizing the animal by placing it at −20 °C for at least 15 min after the experiment. Each body part was then separately weighed using a weighing balance (resolution of 0.1mg; Sartorius BSA2245S-CW, Sartorius AG, Göttingen, Germany).

We digitized the joints/ends of all moving body parts using DeepLabCut, a marker-less pose estimation software (Mathis et al., 2018). In this method, we manually labelled one of every five trials to train a ResNet-50 network with default parameters for 50,000 iterations. For each individual, a separate network was trained and used to analyse all of that individual’s trials recorded under similar experimental conditions. We filtered the position data obtained from DeepLabCut using a third-order low-pass Butterworth filter with a cut-off frequency of 75 Hz to attenuate high-frequency noise arising from the sampling rate and digitization errors between consecutive frames.

To obtain instantaneous CoM speed and trajectory angle, we derived a frame-to-frame displacement vector throughout the video, where the vector magnitude over time represented speed, and its direction represented trajectory/movement angle. From the videos, the take-off time was identified manually as the time instant when the last leg of the animal lost contact with the platform. We labelled the speed and angle corresponding to the time point of take-off as *take-off speed* and *take-off angle*, respectively. Because the mantis could move freely on the take-off platform, its position and hence the perceived target distances and angles varied across trials. Although the mantis estimates the target distance visually, we assume that it adjusts its take-off speed and take-off angle relative to its center of mass. Hence, from the first frame of each video, we measured the target distance and target angle from its eye to the tip of the target platform (Fig. 1C).

### Quantification of peering frequency and amplitude in the variable target distance experiment

To quantify peering kinematics, we separately filmed the lateral head movements of mantises at 50 Hz using a dorsally placed camera (BFS-U3-13Y3M-C, Teledyne Vision Solutions, Ontario, Canada) while varying the target distance. We filmed 3 trials per target distance and manually identified the start and stop of the last peering bout before the mantis jumped towards the intended target. Similar to the previous experiment (Fig. 1A), the distance between platforms was varied by 10 mm or 20 mm, depending on the size of the animal. For each mantis, the distance was varied 4 to 5 times over a total range of 60 to 120 mm. We then digitized the position of the head using DeepLabCut, as previously described. For each trial, the target distance was calculated from the mean head position during the final peering bout to the tip of the target platform. As the mantis was positioned differently on the take-off platform in each trial, the perceived target distance varied across trials. Using the time-series data of lateral head movement, we performed a Fourier transform to extract the dominant frequency and its corresponding amplitude. We restricted the analysis to trials with peering bouts of at least 1 s to provide sufficient temporal sampling for estimating low-frequency components.

### Quantification of the head and body orientation in the variable target angle experiment

To quantify head orientation before jumping, we manually digitised the dorsal and ventral endpoints of the head in the first frame of each video using FIJI (Schindelin et al., 2012) from the high-speed videos of the target angle experiment. Head orientation was defined as the orientation of the vector perpendicular to the line joining the two endpoints of the head (see Fig. 1Ci). To quantify body orientation, we used DeepLabCut to track the anterior and posterior ends of the prothorax throughout the video, as described previously. The orientation of the prothorax with respect to the horizon was used as a proxy for the body orientation (Fig. 1Ci). We compared the target angle with body orientation at three time points: 100 ms before take-off (t= −100 ms), at take-off (t=0), and at touchdown.

### Statistical tests

We quantified the relationships between the independent variables (target distance and target angle) and the dependent variables (take-off speed, take-off angle, head angle, and body angle) by fitting separate simple linear regression models for each individual using ordinary least-squares estimation. This approach allowed us to analyse individual-specific responses to the target parameters. For each regression, we assessed the statistical significance of the relationship by testing the null hypothesis that the slope equals zero against a two-sided alternative. We performed this test using a two-sided Wald test with a Student’s t-distribution. We report the estimated slope (indicating the direction and magnitude of the relationship), the corresponding two-sided p-value, and the coefficient of determination (R-square, representing the proportion of variance in the dependent variable explained by the predictor) for each regression (Supplementary tables 1-10).

## Results

In their arboreal habitats, mantises jump to targets of varying distances and angles, which requires the ability to estimate the distance and angle of the target and generate sufficient force to reach it. We hypothesized that mantises visually estimate the distance and elevational angle of their target using distinct postural movements. Having estimated these, they actively vary both their take-off speed and take-off angle to reach the target. Because the jump trajectories follow the parabolic trajectories of ballistic objects, we modelled the jumps as projectiles on an inclined plane for multiple target distances and angles to predict the take-off speeds and take-off angles necessary to reach the target.

### Modelling the projectile trajectory for fixed target distance and angle

We first determined the range of possibilities available to the mantis modelled as a projectile. Generally, a projectile trajectory is described using the take-off speed (*u*) and take-off angle (*θ*). Because the targets are located at arbitrary distances and angles relative to one another, we derived the generalized equation of projectile motion on an inclined plane (e.g. Crompton and Sellers, 2007) to account for both target distance and angle. This equation requires four input parameters: take-off speed (*u*), take-off angle (*θ*), target distance (*R*) and target angle (*φ*) (Fig 1C), of which (*u, θ*) are modulated by the mantis, whereas (*R, φ*) are fixed by the experimenter.

In a successful jump, the projectile covers the target distance *R* at the angle *φ* by adjusting the take-off speed *u* and take-off angle *θ* as follows (Supplementary material 3):

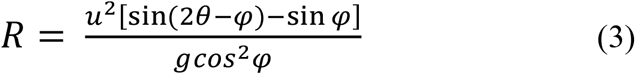

If the projectile starts and ends on level ground (*φ =* 0), we obtain the familiar equation for the projectile range.

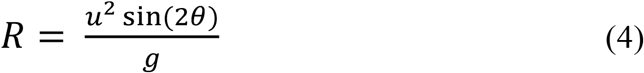

Rearranging (3), we get

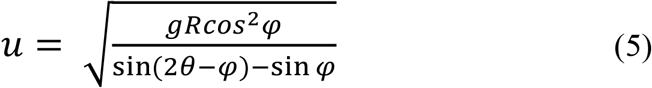

The minimum speed (or energy) to scale the gap requires (derivation in S3),

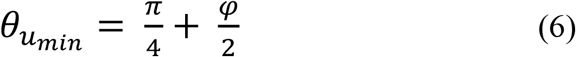

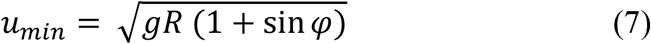

Rearranging (3), we get

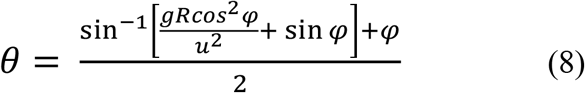

The shortest time required to scale the gap requires the animal to jump at its maximum take-off speed *u*_*max*_, and at an appropriate take-off angle 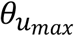,

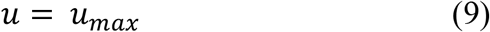

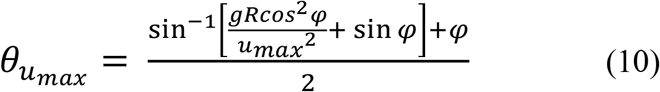

Using these equations, we generated model predictions of the kinematic landscape of take-off speed, take-off angle and target distance for experiments in which the target distance is varied while keeping the target angle constant (Fig. 1F). A similar kinematic landscape was generated for experiments in which the target distance was kept constant while varying the target angle (Fig. 1G). Based on this model, we performed two separate experiments in which either the target angle or the target distance was held constant, and generated predictions for the two jumping strategies.

We additionally explored two distinct strategies that mantises can use to reach a target. In the *speed/energy minimizing strategy*, if mantises minimize their take-off speed (or energy) required to jump, the take-off angle would follow 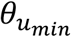 (Eqn. 6). Alternatively, in the *time minimizing strategy*, they minimize the time taken to reach a target, which would require them to jump at a maximum take-off speed of *u*_*max*_ but modulate their take-off angle 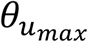 (Eqn. 10).

### Take-off speed and take-off angle increase with target distance

Arboreal habitats are typically discontinuous with varying gaps between substrates. To understand how mantises navigate this habitat, we tested for the effect of target distance on take-off speed and take-off angle. We filmed a total of 439 jumps across 15 individuals while varying the target distance, and quantified their jump kinematics. In all the recorded jumps, the center of mass (CoM) landed close to the nearest edge of the target platform (Fig. 1D), enabling successful grasping of the substrate with their raptorial forelegs (Supplementary movie 1 and 2). Across all trials for an individual, the CoM speed began increasing approximately 75 ms before take-off, as the mantis pushed on its substrate, and attained a maximum value that was proportional to the target distance (Fig. 2A). In contrast, this distance dependent pattern for the CoM angle was less pronounced (Fig. 2B). Across all trails, the take-off speed increased linearly with target distance, with slopes ranging from 2.17 to 5.54 s^−1^ and R^2^ ranging from 0.65 to 0.89 (Fig. 2C; p < 0.05 for all individuals; Wald test; Supplementary Table 1). In most individuals, the take-off angle also increased linearly with target distance with slopes ranging from 44.78 to 348.97 ° m^−1^, but this relationship was somewhat weaker with R^2^ values ranging from 0.16 to 0.83 (Fig. 2D; p < 0.05 in 13 out of 15 individuals; Wald test; Supplementary Table 2). Together, these results indicate that mantises control their jumps by increasing both take-off speed and take-off angle as a function of target distance.

**Figure 2.**
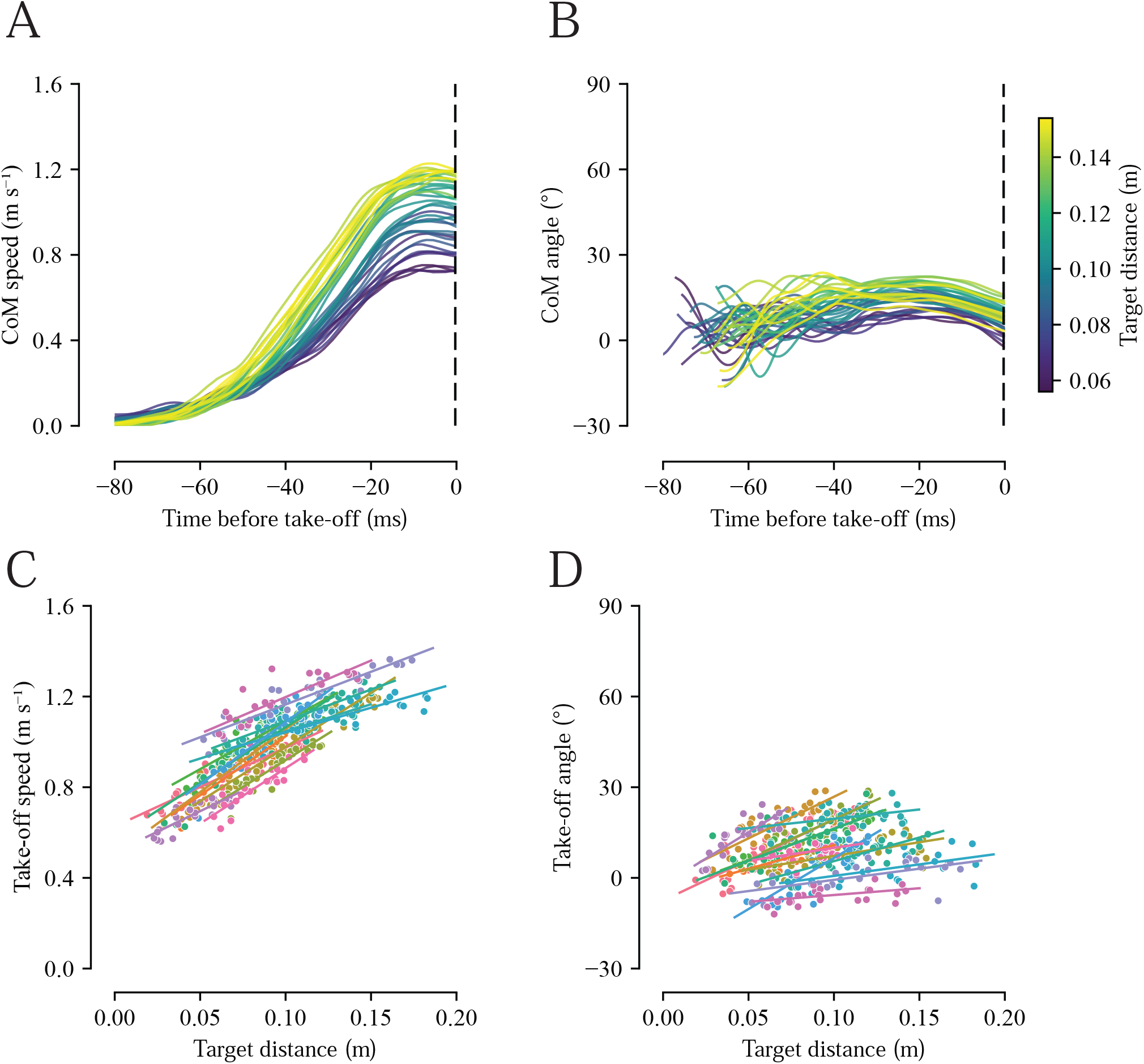
Effect of target distance on take-off speed and angle. Representative traces of an individual’s CoM (A) speed and (B) angle over time before take-off (t = 0, black dashed line) at varying target distances. CoM angle traces are thresholded using a CoM speed of 0.05 ms^−1^ for timeseries visualization. (C-D) Linear regressions of (C) take-off speed and (D) take-off angle as a function of target distance. Data in (C-D) are from a single experiment (15 individuals, 439 trials). Each individual is represented by a distinct color and each trail by a point.

### Peering kinematics correlate with target distance

How do mantises estimate the distance of the target prior to a jump? Locusts and mantises are known to estimate distance using a behavior called ‘peering’, in which insects perform lateral body oscillations before jumping (Sobel, 1990a; Walcher and Kral., 1994; Poteser and Kral, 1995). During peering, insects acquire parallax cues that allow them to estimate distance to the target and accordingly adjust their take-off speed (Sobel, 1990a). We separately filmed a total of 72 jumps from 5 individual mantises from a dorsal perspective and quantified the lateral displacement of the head as they peered toward targets at varying distances. For each mantis, we conducted 15 trials in which we changed the distance based on body size (see methods) and recorded their movements. Across all trials, mantises consistently exhibited peering behavior prior to take-off (e.g., Fig. 3A; Supplementary movie 3). The frequency and amplitude of this peering motion was quantified by applying a Fourier transform to the head position time series (e.g, Fig. 3B). Across trials, the peering frequency decreased with increasing target distance, with slopes ranging from −0.006 to −0.021 Hz m^−1^ and R^2^ ranging from 0.11 to 0.75. (Fig. 3C; p < 0.05 in 4 out of 5 individuals; Wald test; Supplementary Table 3). On the other hand, we did not observe any specific correlation between peering amplitude and target distance (Fig. 3D; p < 0.05 in 1 out of 5 individuals; Wald test; Supplementary Table 4).

**Figure 3.**
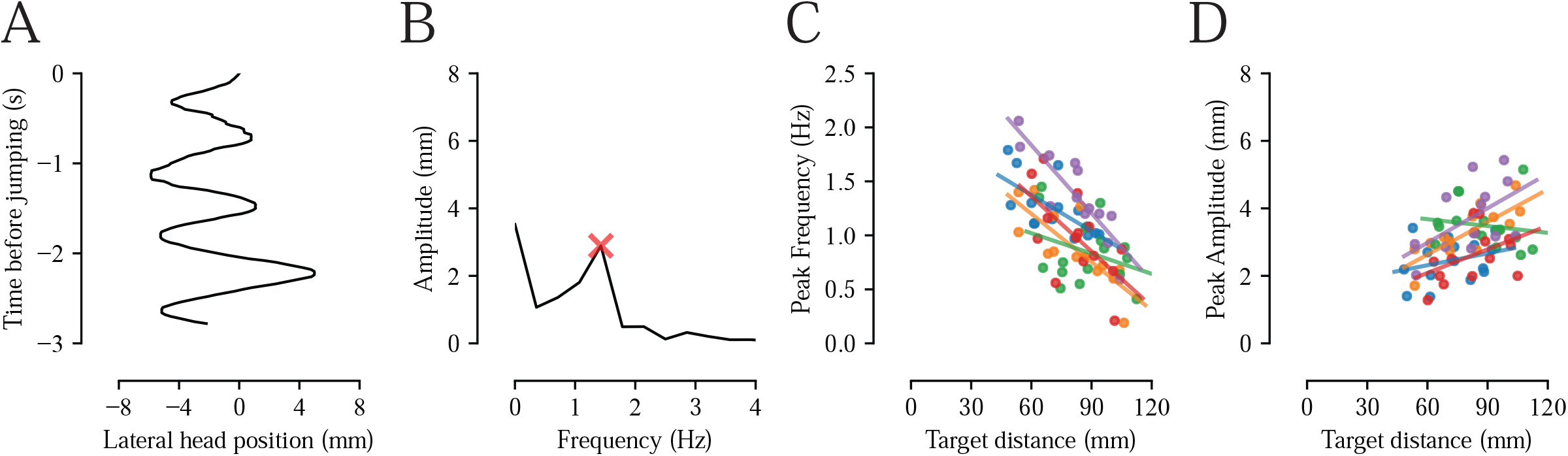
Effect of target distance on peering. (A) Representative trace of the lateral head movement (peering) in a trial. (B) The fast Fourier transform of the trace (A) with the red cross indicating dominant frequency and amplitude. (C-D) Linear regressions of (C) peak frequency and (D) peak amplitude as a function of target distance. Data in (C-D) are from a single experiment (5 individuals and 72 trials). Each individual is represented by a distinct color and each trail by a point.

### Take-off speed decreases and take-off angle increases with the target angle

In the earlier experiments, we varied target distance while keeping target angle constant. However, under natural conditions, mantises jump to targets that vary simultaneously in both distance and angle. To determine how mantises adjust their kinematics when jumping to targets at different target angles, we filmed a total of 241 jumps from 10 individual mantises using a setup in which the target angle was varied from 0° to 90° in steps of 22.5° (Fig. 1B, E; Supplementary movie 4 and 5), with the target distance held constant roughly at 1.5 body lengths for each individual. Across all trials for an individual, the CoM speed began increasing approximately 50 ms before take-off (Fig. 4A) and exhibited a target angle dependent pattern for the CoM angle (Fig. 4B). Across all individuals, take-off speed weakly decreased with increasing target angle, with slopes ranging from −0.0005 to −0.0028 m s^−1^ °^−1^ and R^2^ ranging from 0.05 to 0.86 (Fig. 4C; 8 out of 10 individuals with p < 0.05; Wald test; Supplementary Table 5). We also observed a strong linear relationship between take-off angle and target angle in all individuals, with the regression slopes ranging from 0.59 to 0.83 and R^2^ ranging from 0.91 to 0.98 (Fig. 4D; p < 0.05 for all individuals; Wald test; Supplementary Table 6). Despite the weak effect of target angle on the take-off speeds, mantises primarily reach targets at different angles by systematically modulating their take-off angles.

**Figure 4.**
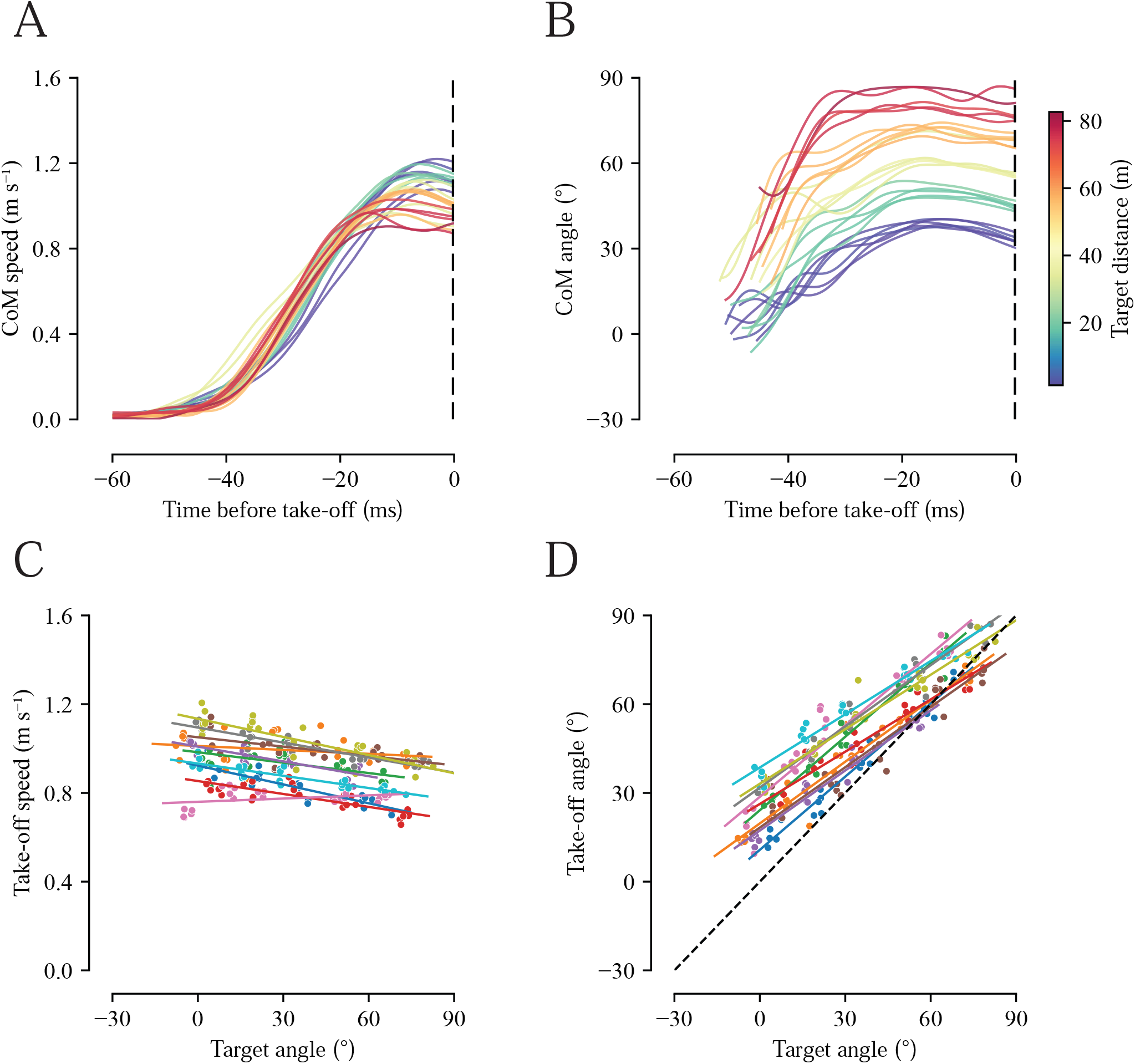
Effect of target angle on take-off speed and angle. Representative traces of an individual’s CoM (A) speed and (B) angle over time before take-off (t = 0, black dashed line) at varying target angles. CoM angle traces are thresholded using a CoM speed of 0.05 ms^−1^ for timeseries visualization. (C-D) Linear regressions of (C) take-off speed and (D) take-off angle as a function of target angle. Data in (C-D) are from a single experiment (10 individuals, 241 trials). Each individual is represented by a distinct color and each trail by a point. The black dashed line in (D) represents a slope of 1.

### Mantises visually assessed the target’s angular position and adjusted their body orientation relative to the target

In the previous experiments, we noticed that the mantises consistently orient their head towards the target before jumping. Quantitative analysis showed that head orientation closely tracked target angle, with slopes ranging from 0.74 to 0.93 and R^2^ ranging from 0.97 to 0.99 (Fig. 5A; p < 0.05 for all individuals; Supplementary Table 7). Mantises also adopted different body postures when jumping to targets at different angles and landed ventral side down on the target (Fig. 5B). We quantified the body angles at three time points – 100 ms before take-off, at take-off and at touchdown. The body angle varied linearly with the target angle at all three stages of the jump - 100 ms before take-off (Fig. 5C; slopes: 0.55 - 0.77; R^2^: 0.93 - 0.98; p < 0.05 for all individuals; Supplementary Table 8), at take-off (Fig. 5D; slopes: 0.67 - 0.90; R^2^: 0.94 - 0.98; p < 0.05 for all individuals; Supplementary Table 9) and at touchdown (Fig. 5E; slopes: 0.68 - 1.45; R^2^: 0.66 - 0.92; p < 0.05 for all individuals; Supplementary Table 10). Together these results suggest that the mantises visually fixate on the target and align their head and body to the target. During the jump, they further reorient their body mid-air to achieve controlled landing on the target ventrally.

**Figure 5.**
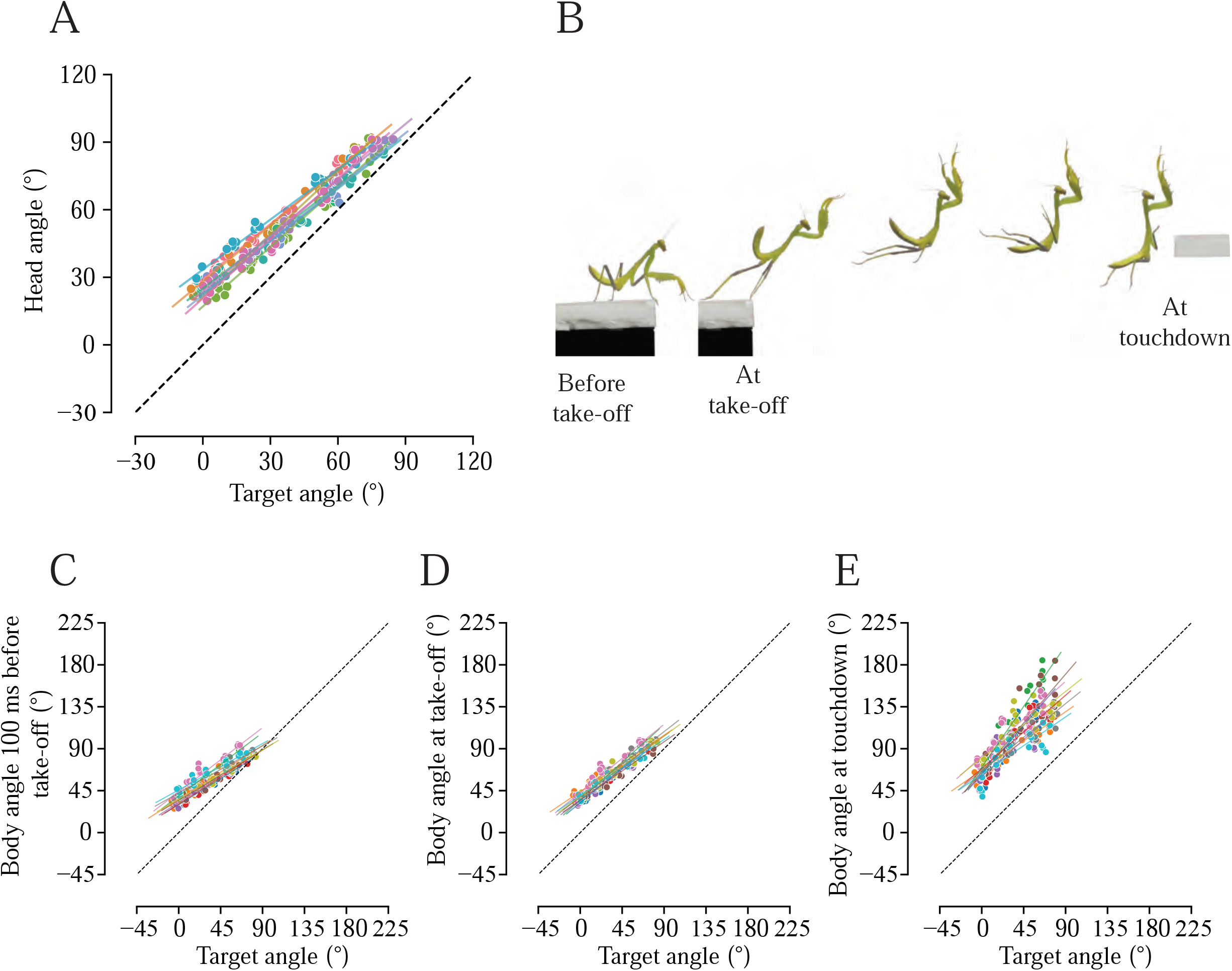
Effect of target angle on head and body orientation. (A) Linear regression of head angle as a function of target angle. (B) Image sequence of mantis nymph during a jump, depicting changes in body posture. (C-E) Linear regressions of body angles at (C) 100ms before take-off, (D) take-off and (E) touchdown as a function of target angle. Data in (A, C-E) are from the same experiment as Fig. 4 (10 individuals, 241 trials). Each individual is represented by a distinct color and each trail by a point. The black dashed line in (D) represents a slope of 1.

### Mantises use flexible strategies rather than optimizing for energy or time

Based on the modelling of the projectiles, we find two strategies: (i) *speed/energy minimizing* and (ii) *time minimizing strategy*. We tested if mantises follow any of the strategies to jump. Because we approximate the mantis as a point particle, we redefine target distance and angle relative to the center of mass at take-off to enable comparison with ideal projectile predictions. (Fig. 6 A-F). In the variable target distance experiment, all mantises increased their take-off speed and take-off angles with target distance (Fig. 6A-C; data for 1 individual shown and the rest in supplementary). In the variable target angle experiment, all the individuals increased their take-off angles, and most individuals mildly decreased take their take-off speeds with target angle (Fig. 6D-F; data for 1 individual shown and the rest in supplementary). The take-off angles were more than half of the target angle. Although the proposed model predicted that mantises should minimize either take-off speed (energy) or time (via constant maximal speed), the observed behavior did not conform to either optimality criterion. This suggests that they use flexible or adaptive strategies when jumping across gaps.

**Figure 6.**
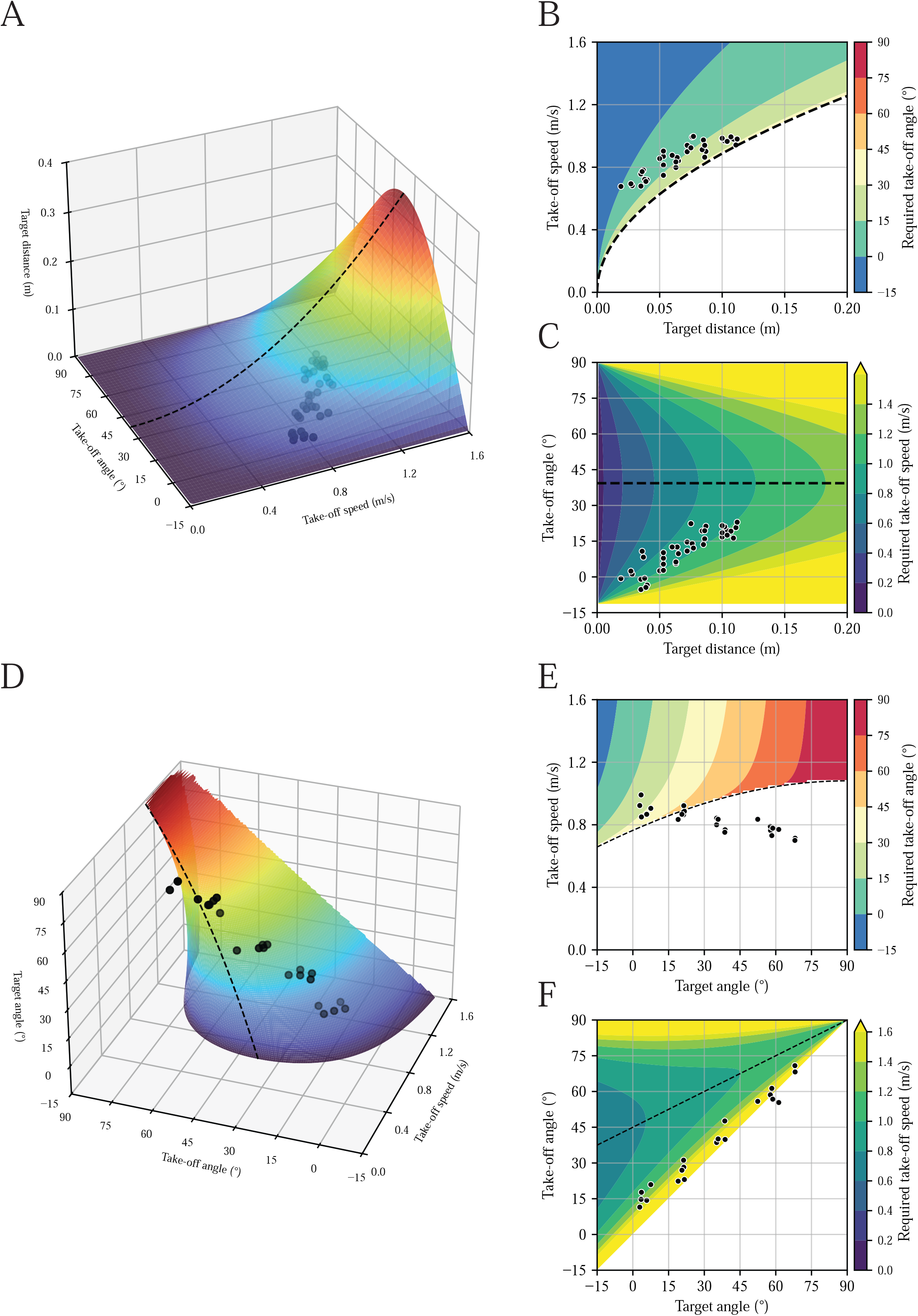
Modelled projectile state-space and comparison with experimental data. (A) 3D state-space of projectile motion with target angle fixed, overlaid with data points from a representative individual (M12). (B-C) Two orthogonal views of the state space showing (B) take-off speed and (C) take-off angle as a function of target distance, with M12 data overlaid. (D) 3D state-space of projectile motion with target distance fixed, overlaid with data points from a representative individual (M25). (E-F) Two orthogonal views of the state space showing (E) take-off speed and (F) take-off angle as a function of target angle, with M25 data overlaid. The black dashed line indicates the take-off *speed/energy minimizing strategy*.

## Discussion

Mantises are diurnal predatory insects that navigate in arboreal environments. Nymphal mantises, which lack wings, frequently rely on jumping to cross gaps exceeding their reach. Using a combination of behavioral assays, high-speed videography and modelling, we investigated how mantises control their targeted jumps. We describe several features of these jumps. First, mantises visually estimate target distance using the peering behavior, similar to locusts and other insects. Second, their take-off speed depends linearly on their target distance. Third, they visually estimate the angle (elevation) of the target and their take-off angle is directly proportional to the target angle. Fourth, they orient their body toward the target prior to take-off and then actively reorient during the jump to achieve accurate landing. Fifth, mantises show flexible jumping strategies rather than relying on any one optimal strategy.

### Role of peering in distance estimation

We report that the peering frequency decreases with an increase in target distance (Fig. 3C) and peering amplitude is uncorrelated with target distance (Fig. 3D). Peering enables distance estimation through position parallax cues, in which distance is estimated as the ratio of the observer’s lateral displacement to the resulting angular displacement of the target, or else through motion parallax cues, in which distance is estimated as the ratio of the observer’s lateral speed to the target’s angular speed across the visual field (Wallace, 1959; Sobel, 1990a; Poteser and Kral, 1995).

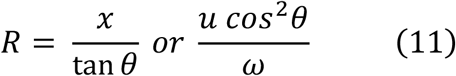

Using small-angle approximation,

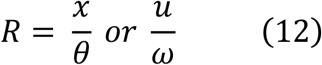

Where *R* is the distance to the target, *x* and *u* are the lateral displacement and speed of the observer, and *θ* and *ω* are the resulting angular displacement and angular speed of the target in the visual field.

For both position and motion parallax, the angular displacement and angular velocity of a target are extracted through visual cues. In contrast, information about the observer’s self-motion may be derived from a combination of proprioceptive signals, motor efferent signals, and visual feedback from self-motion (e.g., optic flow). From behavioral data alone, it is difficult to distinguish whether an animal relies primarily on position parallax or motion parallax to estimate distance, as this requires combining neurophysiological measurements along with behavioral assays.

In principle, monocular cues are sufficient for estimating both absolute and relative distances using parallax or relative motion. With one eye covered and in presence of multiple target substrates, locusts choose the nearest target (Wallace, 1959) and they also jump with higher take-off speeds as compared to binocular conditions (Sobel, 1990a). These observations suggest that binocular vision enhances, but is not strictly required for, distance estimation in locusts. In mantises, however, the evidence remains equivocal: one study reports that mantises perform targeted jumps with one eye occluded (Horridge, 1986), while another study (Walcher and Kral, 1994) could not get such mantises to perform targeted jumps, and hence argues that inputs from both eyes are necessary.

Behavioral studies in the desert locust *Schistocerca gregaria* show that, when the distance to a target is held constant, relative motion between the target and its background influences take-off speed during jumping. This indicates a sensitivity to depth cues that are derived from image motion (Collett and Patterson, 1991). Similarly, in the vinegar fly *Drosophila melanogaster*, gap-crossing decisions depend on perceived distance, and neurons in the optic lobe, including lobula columnar cells, encode motion signals consistent with relative depth estimation (Shomar et al, 2025). These findings suggest that insects can infer spatial layout using motion-based cues, including motion parallax and relative motion, although it is important to demonstrate this in mantises as well.

When multiple objects are present in the visual field, their relative distances can be inferred without explicitly encoding self-motion. In such cases, relative position may be defined as the ratio of angular displacements of two objects during observer movement, whereas relative motion (also referred to as velocity parallax or velocity contrast) corresponds to differences or ratios in their angular velocities across the visual field. In both cases, nearer objects exhibit larger displacements or higher apparent speeds than more distant ones. These cues extend the principles of single-target parallax to multi-object scenes, providing information about relative depth.

Previous studies have reported that peering amplitudes increase with target distance (Collett, 1978; Poteser and Kral, 1995), which is thought to reduce errors in distance estimation since accuracy increases as the observer’s displacement increases when using position based parallax cues. In contrast, our results show that peering amplitude is uncorrelated with target distance. Instead, we find that peering frequency decreases with increasing target distance. This pattern suggests that distance estimation while peering may rely on motion parallax, with a distance dependent modulation of peering frequency rather than amplitude.

Apart from parallax, distance to an object can be estimated visually by binocular disparity, vergence, and stereopsis. Mantises are the only insects reported to use stereopsis to measure depth (Rossel 1983; Nityananda et al., 2016). In experiments in which targets were moved while the insects peered, they mis-estimated the target distance to be further or closer than it was. This resulted in undershooting or overshooting of the jump and missing the target (in locusts - Wallace, 1959; Sobel, 1990a; in mantises - Poteser and Kral, 1995). Again, this suggests that mantises use parallax cues while peering.

### Role of head movements in estimation of elevational angle

Assessing the location of a target substrate requires estimating its distance, the elevational and the azimuthal angle. Mantises rely on head movements to bring objects of interest onto the fovea of the compound eye (Rossel, 1980). Prior to jumping, they orient in the direction of the target, thereby reducing azimuthal error. To estimate elevational angles, they pitch their head up or down until the target is on their fovea (Fig. 1Ci). Thus, head movements, in addition to peering, are required for an accurate estimation of target location. These head movements are sensed by neck proprioceptors, which provide information about head orientation relative to the body. When neck proprioceptors are unilaterally ablated, mantises jump sideways toward a target positioned directly in front of them (Poteser et al., 1998). Thus, proprioceptive feedback may play an important role in estimating target direction. Overall, target localization in mantises likely relies on multisensory integration of visual information (e.g., via peering and foveating) and proprioceptive feedback about head position. The behavioral assay described here is thus useful for probing how insect nervous systems integrate information across sensory modalities.

This study further indicates that mantises estimated target angles since they modulated their take-off angle in relation to the elevation of the target (Fig 4D). In these experiments, because the target was maintained at a constant distance (~1.5 body lengths), distance cues were effectively held constant. Nevertheless, as target elevation increased, take-off speed showed a modest decrease (Fig. 4C). One possible interpretation is that mantises do not fully compensate for gravitational deceleration when executing steeper upward jumps. Under such conditions, successful landing may rely more heavily on the ability of the raptorial forelegs to grasp the target substrate.

### Strategies for jumping in mantises

Ignoring air resistance, the jump of a nymphal praying mantis can be approximated as a projectile trajectory determined by take-off speed and take-off angle. For a target at a fixed distance and elevation, of all the possible combinations of these variables, mantises preferentially select a subset of solutions depending on task demands. From a theoretical perspective, two commonly invoked strategies include a speed/energy-minimizing strategy, which uses the lowest take-off speed required to reach the target, or a time-minimizing strategy, which minimizes time to interception. Saltatorial insects that rely on jumping for routine locomotion (such as grasshoppers) are often assumed to favor energetically efficient trajectories that reduce the cost of transport. In contrast, predators that jump to capture prey (e.g., jumping ants, jumping spiders, praying mantises) may prioritize rapid interception to limit prey escape (Urbani et al., 1994; Hill, 2006; Copeland and Carlson, 1979; Büscher et al., 2026).

Which of these approaches do mantises adopt? To test this, we mapped the state space of feasible take-off speed–angle combinations under ideal projectile motion. Maximum take-off speed is constrained by musculoskeletal properties, including muscle force–velocity characteristics, limb geometry, and body mass. The minimum transit time occurs when the animal jumps at this maximum speed. Lacking direct measurements of individual maximum take-off speeds, we do not impose an explicit constraint in the model, but instead compare observed behavior to an idealized projectile with a defined maximum speed. For a time-minimizing strategy, mantises are expected to jump at a constant maximum speed while adjusting take-off angle (Eqn. 10). In contrast, a speed/energy-minimizing strategy predicts systematic modulation of take-off speed with target distance and elevation, with take-off angles approaching the optimal value of 45° for horizontal targets and shifting with approximately half the target elevation angle (Eqn. 6).

When the target distance was varied, mantises increased both take-off speed and take-off angle with distance (Fig. 6A–C), favouring no specific approach to jumping. When target elevation was varied at a fixed distance, take-off angle increased with target angle, whereas take-off speed decreased modestly (Fig. 6D–F). Although all mantises successfully reached their target, the center of mass did not always coincide with the target at higher elevations. As a result, some data points lie outside the predicted state space. Overall, take-off speed was primarily modulated by target distance, whereas take-off angle was determined by target elevation. These patterns suggest that behavior is flexible and reflects different control rules that lead to distinct regions of the feasible state space. Furthermore, physiological constraints can cause convergence of their predictions at large distances or high elevations, giving the appearance of a strategy shift. Similar interpretations have been proposed in other systems, including jumping spiders (Nabawy et al., 2018) and prosimian primates (Crompton et al., 1993), where energy-efficient trajectories emerge only near performance limits. We treat these strategies as null models against which observed behavior can be evaluated. Mantis performance does not strictly conform to either prediction, underscoring the need to examine behavior across a wide range of target distances and elevations. More generally, inferring locomotor strategies from limited regions of the state space, or interpreting transitions between strategies, requires caution.

### Controlled jumps in insects

A key aim of this study was to understand how insects navigate arboreal habitats, which are structurally complex, three-dimensional environments characterized by discontinuous pathways, variable substrate compliance, and branches differing in width and orientation (Gilman and Irschick, 2013). Navigating in such environments requires traversing gaps via jumping or flight. Unsuccessful attempts, in which the insect misses the target and falls, incur substantial time and energetic costs associated with returning to the canopy. Some arboreal insects, including orchid mantises, stick insects, and certain ant species, mitigate these costs through directed aerial descent following accidental slips, escape jumps, or failed targeted jumps (Yanoviak et al., 2005; Zeng et al., 2015; Zhao et al., 2024). During directed aerial descent, individuals actively steer toward and land on tree trunks rather than falling to the ground, which reduces vertical displacement and improves recovery (Yanoviak et al., 2005). We show here that praying mantises perform controlled, targeted jumps to cross gaps successfully. Such directed jumps represent a widespread mode of locomotion in insects across discontinuous arboreal habitats, which is distinct from other modes of locomotion, such as swimming or flying.

## Acknowledgements

We thank Mr. Abin Ghosh and Ms. Mitali Patil for many discussions and troubleshooting during experiments; Dr. Aashish Satyajith for helping optimize the projectile equations; Mr. Anil Kumar and the Mantis watch group at NCBS, who helped in mantis capture. Funding for this research was provided by the Air Force Office of Scientific Research (AFOSR; grant numbers FA2386-11-1-4057 and FA9550-16-1-0155) and by NCBS, TIFR under project number 12-R&DTFR-5.04-0900.

## Supplementary information legends

**Supplementary Figure 1**. Method to calculate the position of center of mass (CoM) of the animal. (A) Illustration of a mantis nymph in its typical posture before jumping. (B) Digitized joints of each body part using a markerless pose estimation software – DeepLabCut (Mathis et al., 2018). (C) The geometric center of each body part was calculated by taking the mid-point of the two ends of each body part. (D) The position of CoM of the whole body was estimated by taking the average weighted position of each body part.

**Supplementary Movie 1**. High-speed recording of a mantis nymph jumping towards a platform positioned 80 mm away from the take-off platform. The center of mass (CoM) is indicated by an opaque red dot in each frame, with a translucent trail showing its past trajectory. The video was captured at 2000 Hz and played back at 25 Hz.

**Supplementary Movie 2**. High-speed recording of the same mantis nymph (as Supplementary Movie 1) jumping towards a platform positioned 150 mm away from the take-off platform. The CoM is indicated by an opaque red dot in each frame, with a translucent trail showing its past trajectory. The video was captured at 2000 Hz and played back at 25 Hz.

**Supplementary Movie 3**. A top view of a mantis peering before making a jump. The mantis was motivated to peer and jump towards the intended target platform by manually moving a black rod, as shown. The video was captured at 50 Hz and is played back at 50 Hz.

**Supplementary Movie 4**. High-speed recording of a mantis nymph jumping towards a target platform positioned at 0° from the eye of the mantis. The CoM is indicated by an opaque red dot in each frame, with a translucent trail showing its past trajectory. The video was captured at 2000 Hz and played back at 25 Hz

**Supplementary Movie 5**. High-speed recording of the same mantis nymph (as Supplementary Movie 1) jumping towards a target platform positioned 90° from the eye of the mantis. The CoM is indicated by an opaque red dot in each frame, with a translucent trail showing its past trajectory. The video was captured at 2000 Hz and played back at 25 Hz.

**Supplementary Table 1. Linear regression statistics between take-off speed and target distance**. The table reports linear regression summary statistics of take-off speed vs target distance for each individual. The body length, slope, intercept, R-square and standard error are rounded to a decimal point of 2 and the body mass to 3.

**Supplementary Table 2. Linear regression statistics between take-off angle and target distance**. The table reports linear regression summary statistics of take-off angle vs target distance for each individual. The body length, slope, intercept, R-square and standard error are rounded to a decimal point of 2 and the body mass to 3.

**Supplementary Table 3. Linear regression statistics between peering frequency and target distance**. The table reports linear regression summary statistics of peering frequency vs target distance for each individual. The body length, intercept, and R-square are rounded to a decimal point of 2 and the body mass, slope, and standard error to 3.

**Supplementary Table 4. Linear regression statistics between peering amplitude and target distance**. The table reports linear regression summary statistics of peering amplitude vs target distance for each individual. The body length, intercept, and R-square are rounded to a decimal point of 2 and the body mass, slope, and standard error to 3.

**Supplementary Table 5. Linear regression statistics between take-off speed and target angle**. The table reports linear regression summary statistics of take-off speed vs target angle for each individual. The body length, intercept, and R-square are rounded to a decimal point of 2. The body mass is rounded to a decimal point of 3. The slope and standard error to the decimal point of 4.

**Supplementary Table 6. Linear regression statistics between take-off angle and target angle**. The table reports linear regression summary statistics of take-off angle vs target angle for each individual. The body length, slope, intercept, R-square and standard error are rounded to a decimal point of 2 and the body mass to 3.

**Supplementary Table 7. Linear regression statistics between head angle and target angle**. The table reports linear regression summary statistics of head angle vs target angle for each individual. The body length, slope, intercept, R-square and standard error are rounded to a decimal point of 2 and the body mass to 3.

**Supplementary Table 8. Linear regression statistics between body angle 100ms before take-off and target angle**. The table reports linear regression summary statistics of body angle 100ms before take-off vs target angle for each individual. The body length, slope, intercept, R-square and standard error are rounded to a decimal point of 2 and the body mass to 3.

**Supplementary Table 9. Linear regression statistics between body angle at take-off and target angle**. The table reports linear regression summary statistics of body angle at take-off vs target angle for each individual. The body length, slope, intercept, R-square and standard error are rounded to a decimal point of 2 and the body mass to 3.

**Supplementary Table 10. Linear regression statistics between body angle at touchdown and target angle**. The table reports linear regression summary statistics of body angle at touchdown vs target angle for each individual. The body length, slope, intercept, R-square and standard error are rounded to a decimal point of 2 and the body mass to 3.

