## Supplementary material for "The control of targeted jumps in nymphal praying mantises"

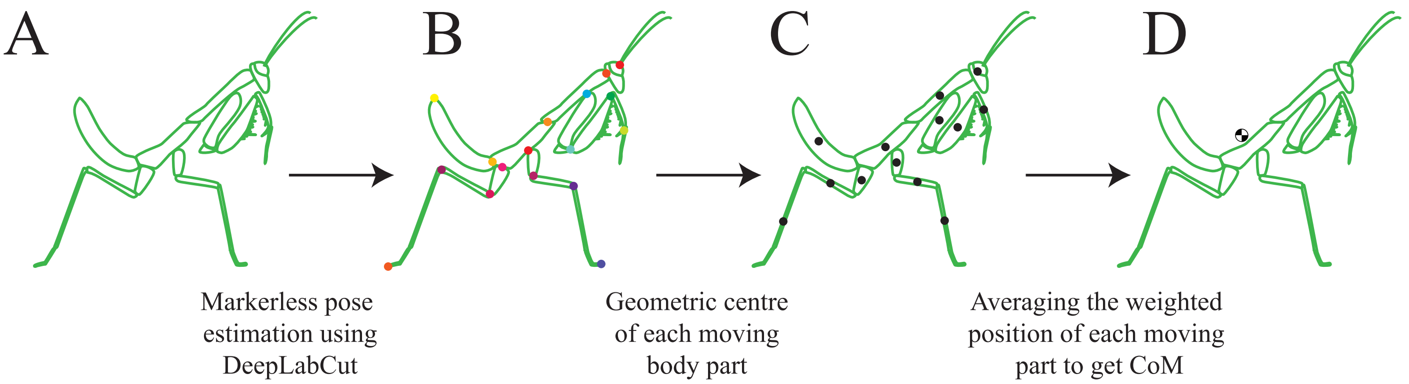
A
B
C
D
Markerless pose estimation using DeepLabCut
Geometric centre of each moving body part
Averaging the weighted position of each moving part to get CoM

### Supplementary Tables 1-10

| Individual ID | Body Length (mm) | Body Mass (g) | Sample size | Slope | Intercept | R-square | p-value | Standard error |
|---|---|---|---|---|---|---|---|---|
| M12 | 27.16 | 0.129 | 40 | 3.48 | 0.63 | 0.81 | 1.89E-15 | 0.27 |
| M13 | 43.65 | 0.625 | 20 | 5.20 | 0.50 | 0.86 | 4.33E-09 | 0.50 |
| M14 | 36.81 | 0.406 | 31 | 5.54 | 0.50 | 0.85 | 2.04E-13 | 0.43 |
| M15 | 61.45 | 1.711 | 40 | 4.78 | 0.50 | 0.89 | 1.11E-19 | 0.28 |
| M16 | 43.66 | 0.686 | 25 | 4.72 | 0.45 | 0.81 | 7.43E-10 | 0.47 |
| M17 | 31.42 | 0.161 | 35 | 4.20 | 0.67 | 0.75 | 1.56E-11 | 0.42 |
| M18 | 36.58 | 0.292 | 35 | 4.90 | 0.57 | 0.81 | 2.26E-13 | 0.42 |
| M19 | 55.09 | 0.782 | 35 | 2.78 | 0.81 | 0.71 | 2.04E-10 | 0.31 |
| M20 | 47.3 | 0.476 | 35 | 2.39 | 0.81 | 0.65 | 5.68E-09 | 0.31 |
| M21 | 64 | 1.297 | 23 | 2.17 | 0.82 | 0.66 | 2.51E-06 | 0.34 |
| M22 | 59.12 | 1.539 | 27 | 5.48 | 0.54 | 0.77 | 2.32E-09 | 0.61 |
| M23 | 70.85 | 0.885 | 25 | 2.89 | 0.88 | 0.88 | 5.18E-12 | 0.22 |
| M24 | 16.31 | 0.02 | 25 | 3.63 | 0.51 | 0.73 | 5.93E-08 | 0.46 |
| M28 | 59.53 | 1.1 | 23 | 3.22 | 0.88 | 0.68 | 6.21E-07 | 0.47 |
| M30 | 61.36 | 1.957 | 20 | 4.89 | 0.40 | 0.76 | 5.66E-07 | 0.65 |

**Supplementary Table 1.** Linear regression statistics for each individual between take-off speed and target distance from Fig. 2C. The body length, slope, intercept, R-square and standard error are rounded to a decimal point of 2 and the body mass to 3.

| Individual ID | Body Length (mm) | Body Mass (g) | Sample size | Slope | Intercept | R-square | p-value | Standard error |
|---|---|---|---|---|---|---|---|---|
| M12 | 27.16 | 0.129 | 40 | 268.17 | -7.50 | 0.78 | 6.89E-14 | 23.41 |
| M13 | 43.65 | 0.625 | 20 | 156.59 | -4.82 | 0.52 | 0.000319 | 35.30 |
| M14 | 36.81 | 0.406 | 31 | 275.13 | -0.67 | 0.64 | 7.07E-08 | 38.44 |
| M15 | 61.45 | 1.711 | 40 | 90.60 | -1.80 | 0.41 | 8.03E-06 | 17.55 |
| M16 | 43.66 | 0.686 | 25 | 259.76 | -6.64 | 0.69 | 2.39E-07 | 35.99 |
| M17 | 31.42 | 0.161 | 35 | 207.71 | -4.54 | 0.83 | 3.15E-14 | 16.39 |
| M18 | 36.58 | 0.292 | 35 | 212.33 | -4.97 | 0.61 | 3.44E-08 | 29.71 |
| M19 | 55.09 | 0.782 | 35 | 155.95 | -10.04 | 0.57 | 1.35E-07 | 23.37 |
| M20 | 47.3 | 0.476 | 35 | 61.33 | 13.43 | 0.20 | 0.007436 | 21.50 |
| M21 | 64 | 1.297 | 23 | 76.18 | -6.97 | 0.25 | 0.016315 | 29.18 |
| M22 | 59.12 | 1.539 | 27 | 337.66 | -27.20 | 0.72 | 2.06E-08 | 41.90 |
| M23 | 70.85 | 0.885 | 25 | 73.96 | -8.06 | 0.38 | 0.001019 | 19.67 |
| M24 | 16.31 | 0.02 | 25 | 348.97 | -2.35 | 0.71 | 1.12E-07 | 46.17 |
| M28 | 59.53 | 1.1 | 23 | 44.78 | -10.15 | 0.16 | 0.05577 | 22.17 |
| M30 | 61.36 | 1.957 | 20 | 85.57 | 1.37 | 0.19 | 0.055788 | 41.85 |

**Supplementary Table 2.** Linear regression statistics for each individual between take-off angle and target distance from Fig. 2D. The body length, slope, intercept, R-square and standard error are rounded to a decimal point of 2 and the body mass to 3.

| Individual ID | Body Length (mm) | Body Mass (g) | Sample size | Slope | Intercept | R-square | p-value | Standard error |
|---|---|---|---|---|---|---|---|---|
| M43 | 27.74 | 0.148 | 15 | -0.011 | 2.04 | 0.44 | 6.98E-03 | 0.003 |
| M44 | 50.08 | 0.710 | 14 | -0.015 | 2.09 | 0.62 | 8.66E-04 | 0.003 |
| M46 | 29.57 | 0.106 | 16 | -0.006 | 1.40 | 0.11 | 2.16E-01 | 0.005 |
| M47 | 61.38 | 0.529 | 14 | -0.017 | 2.38 | 0.39 | 1.66E-02 | 0.006 |
| M48 | 64.61 | 0.317 | 13 | -0.021 | 3.10 | 0.75 | 1.44E-04 | 0.004 |

**Supplementary Table 3**. Linear regression statistics for each individual between peering frequency and target distance from Fig. 3C. The body length, intercept, and R-square are rounded to a decimal point of 2 and the body mass, slope and standard error to 3.

| Individual ID | Body Length (mm) | Body Mass (g) | Sample size | Slope | Intercept | R-square | p-value | Standard error |
|---|---|---|---|---|---|---|---|---|
| M43 | 27.74 | 0.148 | 15 | 0.012 | 1.61 | 0.08 | 3.02E-01 | 0.011 |
| M44 | 50.08 | 0.710 | 14 | 0.032 | 0.69 | 0.62 | 8.57E-04 | 0.007 |
| M46 | 29.57 | 0.106 | 16 | -0.007 | 4.08 | 0.02 | 5.63E-01 | 0.011 |
| M47 | 61.38 | 0.529 | 14 | 0.023 | 0.68 | 0.20 | 1.12E-01 | 0.014 |
| M48 | 64.61 | 0.317 | 13 | 0.034 | 0.93 | 0.28 | 6.24E-02 | 0.016 |

**Supplementary Table 4**. Linear regression statistics for each individual between peering amplitude and target distance from Fig. 3D. The body length, intercept, and R-square are rounded to a decimal point of 2 and the body mass, slope and standard error to 3

| Individual ID | Body length (mm) | Body mass (g) | Sample size | Slope | Intercept | R-square | p-value | Standard error |
|---|---|---|---|---|---|---|---|---|
| M25 | 27.16 | 0.129 | 20 | -0.0028 | 0.92 | 0.74 | 1.56E-07 | 0.0004 |
| M26 | 43.65 | 0.625 | 25 | -0.0006 | 1.01 | 0.05 | 0.259856 | 0.0005 |
| M27 | 36.81 | 0.406 | 25 | -0.0016 | 0.98 | 0.32 | 0.00344 | 0.0005 |
| M29 | 61.45 | 1.711 | 25 | -0.0019 | 0.85 | 0.65 | 1.11E-06 | 0.0003 |
| M31 | 43.66 | 0.686 | 20 | -0.0022 | 1.01 | 0.72 | 2.28E-06 | 0.0003 |
| M32 | 31.42 | 0.161 | 25 | -0.0014 | 1.05 | 0.39 | 0.000844 | 0.0004 |
| M35 | 36.58 | 0.292 | 25 | 0.0005 | 0.76 | 0.10 | 0.129628 | 0.0003 |
| M36 | 55.09 | 0.782 | 25 | -0.0023 | 1.09 | 0.86 | 3.18E-11 | 0.0002 |
| M37 | 47.3 | 0.476 | 26 | -0.0027 | 1.13 | 0.66 | 4.54E-07 | 0.0004 |
| M39 | 64 | 1.297 | 25 | -0.0018 | 0.93 | 0.78 | 5.15E-09 | 0.0002 |

**Supplementary Table 5.** Linear regression statistics for each individual between take-off speed and target angle from Fig. 4C. The body length, intercept, and R-square are rounded to a decimal point of 2. The body mass is rounded to a decimal point of 3. The slope and standard error to the decimal point of 4.

| Individual ID | Body length (mm) | Body mass (g) | Sample size | Slope | Intercept | R-square | p-value | Standard error |
|---|---|---|---|---|---|---|---|---|
| M25 | 27.16 | 0.129 | 20 | 0.82 | 10.77 | 0.97 | 3.38E-18 | 0.03 |
| M26 | 43.65 | 0.625 | 25 | 0.70 | 19.69 | 0.95 | 4.18E-16 | 0.03 |
| M27 | 36.81 | 0.406 | 25 | 0.83 | 24.14 | 0.98 | 5.69E-20 | 0.03 |
| M29 | 61.45 | 1.711 | 25 | 0.59 | 26.27 | 0.91 | 1.80E-13 | 0.04 |
| M31 | 43.66 | 0.686 | 20 | 0.68 | 17.28 | 0.96 | 8.18E-14 | 0.03 |
| M32 | 31.42 | 0.161 | 25 | 0.68 | 18.29 | 0.92 | 3.99E-14 | 0.04 |
| M35 | 36.58 | 0.292 | 25 | 0.80 | 28.68 | 0.92 | 7.02E-14 | 0.05 |
| M36 | 55.09 | 0.782 | 25 | 0.69 | 31.80 | 0.97 | 9.23E-19 | 0.03 |
| M37 | 47.3 | 0.476 | 26 | 0.61 | 33.10 | 0.95 | 1.50E-17 | 0.03 |
| M39 | 64 | 1.297 | 25 | 0.60 | 38.60 | 0.95 | 2.68E-16 | 0.03 |

**Supplementary Table 6.** Linear regression statistics for each individual between take-off angle and target angle from Fig. 4D. The body length, slope, intercept, R-square and standard error are rounded to a decimal point of 2 and the body mass to 3.

| Individual ID | Body length (mm) | Body mass (g) | Sample size | Slope | Intercept | R-square | p-value | Standard error |
|---|---|---|---|---|---|---|---|---|
| M25 | 27.16 | 0.129 | 20 | 0.81 | 29.22 | 0.98 | 2.78E-18 | 0.02 |
| M26 | 43.65 | 0.625 | 25 | 0.83 | 28.13 | 0.99 | 4.31E-23 | 0.02 |
| M27 | 36.81 | 0.406 | 25 | 0.93 | 21.21 | 0.98 | 4.48E-20 | 0.03 |
| M29 | 61.45 | 1.711 | 25 | 0.88 | 17.04 | 0.97 | 1.66E-19 | 0.03 |
| M31 | 43.66 | 0.686 | 20 | 0.76 | 25.00 | 0.97 | 2.44E-15 | 0.03 |
| M32 | 31.42 | 0.161 | 25 | 0.78 | 22.63 | 0.98 | 3.70E-21 | 0.02 |
| M35 | 36.58 | 0.292 | 25 | 0.74 | 33.43 | 0.99 | 3.99E-23 | 0.02 |
| M36 | 55.09 | 0.782 | 25 | 0.80 | 22.11 | 0.98 | 1.69E-21 | 0.02 |
| M37 | 47.3 | 0.476 | 26 | 0.83 | 23.71 | 0.99 | 3.69E-26 | 0.02 |
| M39 | 64 | 1.297 | 25 | 0.89 | 20.63 | 0.97 | 2.25E-19 | 0.03 |

**Supplementary Table 7.** Linear regression statistics for each individual between head angle and target angle from Fig. 5A. The body length, slope, intercept, R-square and standard error are rounded to a decimal point of 2 and the body mass to 3.

| Individual ID | Body length (mm) | Body mass (g) | Sample size | Slope | Intercept | R-square | p-value | Standard error |
|---|---|---|---|---|---|---|---|---|
| M25 | 27.16 | 0.129 | 20 | 0.62 | 30.05 | 0.96 | 3.17E-15 | 0.03 |
| M26 | 43.65 | 0.625 | 25 | 0.64 | 36.12 | 0.96 | 1.93E-17 | 0.03 |
| M27 | 36.81 | 0.406 | 25 | 0.75 | 39.24 | 0.96 | 2.18E-17 | 0.03 |
| M29 | 61.45 | 1.711 | 25 | 0.64 | 30.69 | 0.95 | 2.35E-16 | 0.03 |
| M31 | 43.66 | 0.686 | 20 | 0.65 | 32.43 | 0.94 | 8.80E-13 | 0.04 |
| M32 | 31.42 | 0.161 | 25 | 0.60 | 34.62 | 0.96 | 3.13E-17 | 0.03 |
| M35 | 36.58 | 0.292 | 25 | 0.77 | 43.66 | 0.94 | 7.53E-16 | 0.04 |
| M36 | 55.09 | 0.782 | 25 | 0.56 | 42.51 | 0.97 | 6.95E-19 | 0.02 |
| M37 | 47.3 | 0.476 | 26 | 0.59 | 36.11 | 0.98 | 9.27E-21 | 0.02 |
| M39 | 64 | 1.297 | 25 | 0.55 | 45.78 | 0.93 | 9.49E-15 | 0.03 |

**Supplementary Table 8.** Linear regression statistics for each individual between body angle 100ms before take-off and target angle from Fig. 5C. The body length, slope, intercept, R-square and standard error are rounded to a decimal point of 2 and the body mass to 3.

| Individual ID | Body length (mm) | Body mass (g) | Sample size | Slope | Intercept | R-square | p-value | Standard error |
|---|---|---|---|---|---|---|---|---|
| M25 | 27.16 | 0.129 | 20 | 0.82 | 32.05 | 0.97 | 2.02E-15 | 0.04 |
| M26 | 43.65 | 0.625 | 25 | 0.67 | 43.95 | 0.96 | 6.68E-18 | 0.03 |
| M27 | 36.81 | 0.406 | 25 | 0.90 | 35.08 | 0.97 | 1.72E-19 | 0.03 |
| M29 | 61.45 | 1.711 | 25 | 0.73 | 37.90 | 0.96 | 1.18E-17 | 0.03 |
| M31 | 43.66 | 0.686 | 20 | 0.74 | 34.68 | 0.95 | 5.98E-13 | 0.04 |
| M32 | 31.42 | 0.161 | 25 | 0.70 | 34.80 | 0.98 | 3.31E-20 | 0.02 |
| M35 | 36.58 | 0.292 | 25 | 0.81 | 42.99 | 0.94 | 6.19E-16 | 0.04 |
| M36 | 55.09 | 0.782 | 25 | 0.78 | 38.42 | 0.98 | 2.09E-21 | 0.02 |
| M37 | 47.3 | 0.476 | 26 | 0.72 | 38.78 | 0.96 | 8.23E-19 | 0.03 |
| M39 | 64 | 1.297 | 25 | 0.67 | 39.66 | 0.96 | 1.08E-17 | 0.03 |

**Supplementary Table 9.** Linear regression statistics for each individual between body angle at take-off and target angle from Fig. 5D. The body length, slope, intercept, R-square and standard error are rounded to a decimal point of 2 and the body mass to 3.

| Individual ID | Body length (mm) | Body mass (g) | Sample size | Slope | Intercept | R-square | p-value | Standard error |
|---|---|---|---|---|---|---|---|---|
| M25 | 27.16 | 0.129 | 20 | 1.16 | 62.74 | 0.82 | 1.67E-08 | 0.12 |
| M26 | 43.65 | 0.625 | 25 | 0.69 | 67.12 | 0.86 | 3.65E-11 | 0.06 |
| M27 | 36.81 | 0.406 | 25 | 1.45 | 70.02 | 0.92 | 6.72E-14 | 0.09 |
| M29 | 61.45 | 1.711 | 25 | 0.87 | 68.74 | 0.80 | 1.42E-09 | 0.09 |
| M31 | 43.66 | 0.686 | 20 | 0.88 | 60.31 | 0.66 | 1.31E-05 | 0.15 |
| M32 | 31.42 | 0.161 | 25 | 1.17 | 65.03 | 0.77 | 6.73E-09 | 0.13 |
| M35 | 36.58 | 0.292 | 25 | 0.89 | 81.78 | 0.81 | 7.68E-10 | 0.09 |
| M36 | 55.09 | 0.782 | 25 | 0.80 | 66.04 | 0.87 | 9.00E-12 | 0.06 |
| M37 | 47.3 | 0.476 | 26 | 0.79 | 79.95 | 0.74 | 2.02E-08 | 0.10 |
| M39 | 64 | 1.297 | 25 | 0.68 | 62.50 | 0.66 | 7.39E-07 | 0.10 |

**Supplementary Table 10.** Linear regression statistics for each individual between body angle at touchdown and target angle from Fig. 5E. The body length, slope, intercept, R-square and standard error are rounded to a decimal point of 2 and the body mass to 3.

### Supplementary material 1

#### Deriving the equation for the range of a projectile on an inclined plane

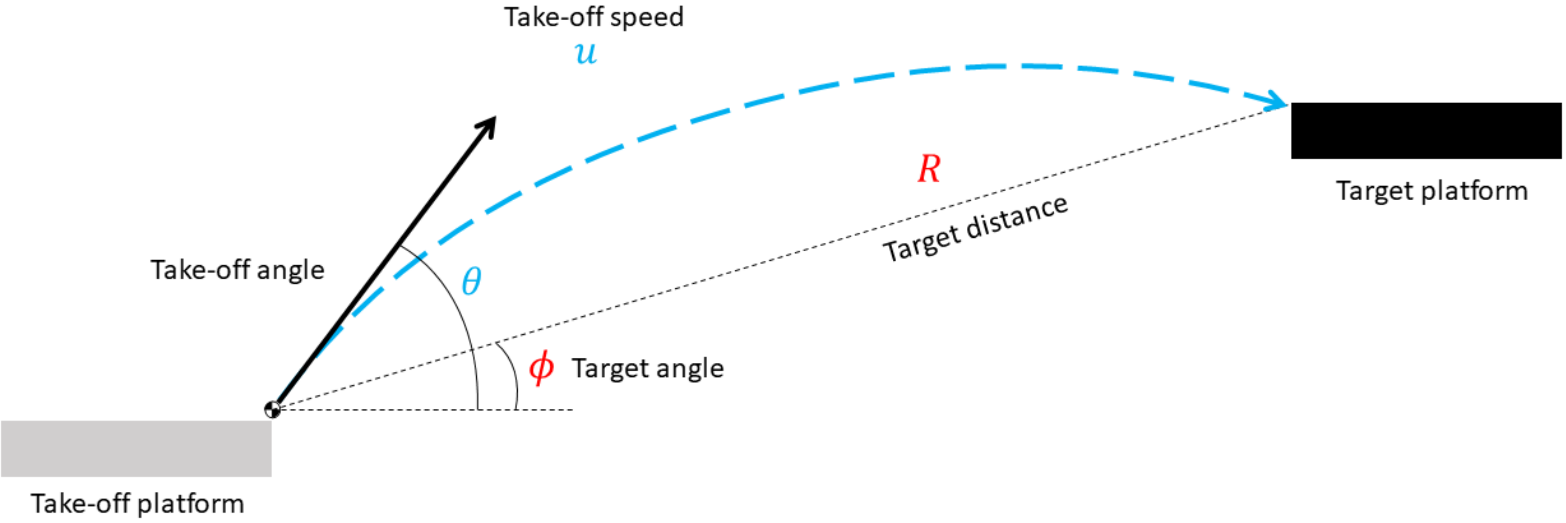


Figure 1: Projectile motion on an inclined plane

Given the following kinematic equations:

$$
\begin{aligned}
v &= u + at \\
s &= ut + \frac{1}{2}at^2 \qquad (1) \\
v^2 &= u^2 + 2as \\
s &= \frac{(u+v)}{2}t
\end{aligned}
$$

Only equation (1) can be used since $v$ is unknown.

Known parameters are $u$, $\theta$, $R$, and $\phi$
In the horizontal direction, from equation (1):
a = 0 in the horizontal direction,

$$R\cos\phi = u\cos\theta t \tag{2}$$

$$\boxed{t = \frac{R\cos\phi}{u\cos\theta}} \tag{3}$$

In the vertical direction, from equation (1):

$$R\sin\phi = u\sin\theta t - \frac{1}{2}gt^2 \tag{4}$$

Solving for $t$ by dividing equation (4) by equation (2):

$$\frac{R\sin\phi}{R\cos\phi} = \frac{u\sin\theta t - \frac{1}{2}gt^2}{u\cos\theta t}$$

$$\tan\phi = \tan\theta - \frac{gt}{2u\cos\theta}$$

Rearranging to get $t$:

$$t = \frac{2u\cos\theta}{g}\left(\tan\theta - \tan\phi\right) \tag{5}$$

$$\boxed{t = \frac{2u\sin(\theta-\phi)}{g\cos\phi}} \tag{6}$$

Substituting $t$ from equation (3) into equation (4):

$$R\sin\phi = u\sin\theta\frac{R\cos\phi}{u\cos\theta} - \frac{1}{2}g\left(\frac{R\cos\phi}{u\cos\theta}\right)^2$$

Multiplying both sides by $\frac{2\cos^2\theta}{R}$:

$$2\sin\phi\cos^2\theta = 2\sin\theta\cos\theta\cos\phi - \frac{gR\cos^2\phi}{u^2}$$

Since $2\sin\theta\cos\theta = \sin 2\theta$ and $\cos^2\theta = \frac{1+\cos 2\theta}{2}$:

$$2\sin\phi\left(\frac{1+\cos 2\theta}{2}\right) = \sin 2\theta\cos\phi - \frac{gR\cos^2\phi}{u^2}$$

$$\sin\phi + \sin\phi\cos 2\theta = \sin 2\theta\cos\phi - \frac{gR\cos^2\phi}{u^2}$$

$$\frac{gR\cos^2\phi}{u^2} = \sin 2\theta\cos\phi - \cos 2\theta\sin\phi - \sin\phi$$

Since $\sin A\cos B - \cos A\sin B = \sin(A-B)$:

$$\frac{gR\cos^2\phi}{u^2} = \sin(2\theta-\phi) - \sin\phi$$

Finally, solving for $R$:

$$\boxed{R = \frac{u^2(\sin(2\theta-\phi) - \sin\phi)}{g\cos^2\phi}} \tag{7}$$

Rearranging equation (7) to solve for $u$:

$$u^2 = \frac{gR\cos^2\phi}{\sin(2\theta-\phi) - \sin\phi} \tag{8}$$

$$\boxed{u = \sqrt{\frac{gR\cos^2\phi}{\sin(2\theta - \phi) - \sin\phi}}} \tag{9}$$

Rearranging equation (7) to solve for $\theta$:

$$\sin(2\theta - \phi) - \sin\phi = \frac{gR\cos^2\phi}{u^2}$$

$$\sin(2\theta - \phi) = \frac{gR\cos^2\phi}{u^2} + \sin\phi$$

$$2\theta - \phi = \sin^{-1}\left(\frac{gR\cos^2\phi}{u^2} + \sin\phi\right)$$

$$\boxed{\theta = \frac{\sin^{-1}\left(\frac{gR\cos^2\phi}{u^2} + \sin\phi\right) + \phi}{2}} \tag{10}$$

To find the landing speed $v$ of the projectile,
from equation

$$v^2 = u^2 + 2as$$

$$v_x^2 = (u\cos\theta)^2$$

$$v_y^2 = (u\sin\theta)^2 - 2gR\sin\phi$$

By geometry

$$v = \sqrt{v_x^2 + v_y^2}$$

$$v = \sqrt{(u\cos\theta)^2 + (u\sin\theta)^2 - 2gR\sin\phi}$$

$$\boxed{v = \sqrt{u^2 - 2gR\sin\phi}} \tag{11}$$

To find the landing angle $\alpha$, from the equation

$$v = u + at$$

In the horizontal direction,

$$v_x = u\cos\theta$$

In the vertical direction,

$$v_y = u\sin\theta - gt$$

Substituting t from equation (5) we get

$$v_y = u\sin\theta - 2u\cos\theta(\tan\theta - \tan\phi))$$

By geometry

$$\tan\alpha = \left(\frac{v_y}{v_x}\right)$$

Substituting $v_x$ and $v_y$ from equations ...., we get

$$\tan\alpha = \frac{u\sin\theta - 2u\cos\theta(\tan\theta - \tan\phi)}{u\cos\theta}$$

$$\tan\alpha = \tan\theta - 2\tan\theta + 2\tan\phi$$

$$\tan\alpha = 2\tan\phi - \tan\theta$$

$$\boxed{\alpha = \tan^{-1}(2\tan\phi - \tan\theta)} \tag{11}$$

#### Minimising take-off speed for a given target distance and target angle

From equation (9), the take-off speed is given by:

$$u = \sqrt{\frac{gR\cos^2\phi}{\sin(2\theta - \phi) - \sin\phi}}$$

Differentiating this with respect to $\theta$:
$g$, $R$, $\phi$ are constants,

$$\frac{du}{d\theta} = \sqrt{gR\cos^2\phi}\,(-\cos(2\theta - \phi))\left(\frac{1}{\sin(2\theta - \phi) - \sin\phi}\right)^{\frac{3}{2}} \tag{13}$$

Equating equation (13) to 0:

$$\sqrt{gR\cos^2\phi}\,(-\cos(2\theta - \phi))\left(\frac{-1}{\sin\phi - \sin(2\theta - \phi)}\right)^{\frac{3}{2}} = 0$$

Solving for $\theta$:

$$\cos(2\theta - \phi) = 0$$

$$2\theta - \phi = \cos^{-1}(0)$$

$$2\theta - \phi = \frac{\pi}{2} \tag{14}$$

$$2\theta = \frac{\pi}{2} + \phi$$

$$\theta = \frac{\pi}{4} + \frac{\phi}{2} \tag{15}$$

Differentiating equation (13) with respect to $\theta$ again:

$$\frac{d^2u}{d\theta^2} = \sqrt{gR\cos^2\phi}\sqrt{\frac{-1}{\sin\phi - \sin(2\theta - \phi)}}\left(\frac{(3\cos^2(2\theta - \phi) - 2\sin(2\theta - \phi)(\sin\phi - \sin(2\theta - \phi)))}{(\sin\phi - \sin(2\theta - \phi))^2}\right) \tag{16}$$

Substituting the value of $\theta$ from equation (15) or the value of $2\theta - \phi$ from equation (14) into equation (16):

$$\frac{d^2u}{d\theta^2} = \sqrt{gR\cos^2\phi}\sqrt{\frac{1}{1 - \sin\phi}}\frac{-2(\sin\phi - 1)}{(\sin\phi - 1)^2}$$

$$\frac{d^2u}{d\theta^2} = \sqrt{gR\cos^2\phi}\frac{2\sqrt{1 - \sin\phi}}{(\sin\phi - 1)^2}$$

Since $\frac{d^2u}{d\theta^2}$ is positive, the value of $\theta$ is $\frac{\pi}{4} + \frac{\phi}{2}$ at the minima of take-off speed, $u$.

$$\boxed{u_{\min} = \sqrt{gR\,(1 + \sin\phi)} \quad \text{when} \quad \theta = \frac{\pi}{4} + \frac{\phi}{2}}$$
